# The Occurrence Birth-Death Diffusion Process: Unraveling Diversification Histories with Fossils and Heterogeneous Rates

**DOI:** 10.1101/2025.08.26.672414

**Authors:** Jérémy Andréoletti, Ignacio Quintero, Hélène Morlon

**Author notes:** Corresponding authors, 39 avenue François-Vincent Raspail, 94110 Arcueil, France, AND, 46 rue d’Ulm, 75005 Paris.

## Abstract

Phylogenetic diversification analyses often overlook the wealth of information contained in fossils that cannot be directly integrated into phylogenetic trees (called fossil occurrences), as well as fine-grained variations in speciation and extinction rates across lineages. Addressing both of these shortcomings, we develop the Occurrence Birth-Death Diffusion (OBDD) process. This process incorporates the sampling of fossil occurrences by extending the recently developed Fossilized Birth-Death Diffusion (FBDD) process, thereby taking advantage of a clade’s entire fossil record to infer its evolutionary history. To mitigate the computational challenges associated with the OBDD process, we employ Bayesian data augmentation methods. We illustrate our approach with an empirical analysis of cetacean diversification. Our analyses show that the integration of fossils is essential to recover diversification rates and richness trajectories. More specifically, all types of fossils are highly informative about past diversification dynamics, but fossil occurrences do not provide significant additional information when the phylogeny already contains many fossils. We make the OBDD available in the Julia package Tapestree.

## Introduction

The study of diversification histories sheds light on the patterns and processes that drive the emergence, extinction, and evolution of species. Initiated by extensive paleontological research on the fossil record, the field has seen significant advancement with the introduction and refinement of phylogenetic diversification models. These models represent diversification as a birth-death (BD) process, where each lineage can give birth or go extinct at given rates (commonly denoted as *λ* for speciation and *μ* for extinction) (Nee et al., 1994; Stadler, 2009, 2010). Early iterations of these models were developed for phylogenies of extant-only taxa and assumed constant rates over time and across lineages, but they have progressively incorporated more complexity, e.g. allowing for temporal and among-lineage variation in rates and dependencies on traits or environmental factors (Stadler, 2013; Morlon, 2014; Morlon et al., 2024).

Recently, substantial progress has been made in integrating direct fossil evidence into phylogenetic diversification analyses by adding sampling processes to the birth-death process, thus leveraging the extensive information contained in paleontological databases. This resulted in the development of the Fossilized Birth-Death (FBD) model, which integrates into a phylogeny fossils with adequate morphological or taxonomic information (Stadler, 2010; Heath et al., 2014). However, this information is not available for most fossils, leading traditional FBD models to ignore a large portion of the fossil record. In fact, due to the effort required to compile these data, researchers often include only one fossil per morphospecies in matrices of morphological characters (Wright et al., 2022). In response, the Occurrence Birth-Death (OBD) model was developed to include fossil occurrences, that is, the remaining fossils that lack the publicly compiled morphological or taxonomic information that would allow them to be placed in a phylogeny (Vaughan et al., 2019; Gupta et al., 2020; Andréoletti et al., 2022). From now on, we will use “fossil” or “phylogenetic fossil” to designate those in the tree and “occurrence” or “fossil occurrence” for those that are not. Although fossil occurrences are not as informative as phylogenetic fossils, variation in their number over time can reflect variation in a clade’s total species richness and inform the underlying speciation and extinction rates. This is analogous to the complementary information provided by genome sequences (analogous to fossil characters) and incidence data (analogous to occurrences) to understand pathogen dynamics, and, in fact, the same phylodynamic tools are used in this context (MacPherson et al., 2022; Stadler et al., 2024). Analyses of the OBD model in the context of pathogen phylodynamics have shown that sampling times (incidence data) often play a key role in driving inference when sequence diversity is low; in contrast, in scenarios of rapid evolution, sequence data provide the dominant source of information (Featherstone et al., 2023). This suggests that occurrences will be particularly useful when the placement of fossils in the phylogeny is uncertain, for example due to few or similar morphological characters. The OBD model was first implemented by Vaughan et al. (2019) in the phylogenetic software BEAST2 (Bouckaert et al., 2019) with a particle filter algorithm. Next, Gupta et al. (2020) developed a faster algorithm and Andréoletti et al. (2022) extended the model to account for the piecewise constant rate variation and implemented it in RevBayes (Höhna et al., 2016). Both implementations allow full joint inference of phylogenetic trees and diversification (or epidemiological) parameters. However, their application to large trees is hampered by computational limitations, primarily due to a numerically intensive integration step required for the likelihood computation. A fast approximation of this model using a method called TimTam (Zarebski et al., 2025) has been implemented in Beast2 but is currently limited to a scenario with lineage removal at sampling, which makes sense in the case of sampling pathogen genomes, but is not realistic in the case of sampling fossils. In addition, all current implementations of the OBD model assume homogeneous diversification rates across lineages.

Substantial progress has also been made in developing diversification models that accommodate rate heterogeneity, in which rates can differ between portions of the tree, either due to rare clade-wide shifts (Rabosky, 2014; Mitchell et al., 2019; Maddison et al., 2007; Caetano et al., 2018), or more gradual branch-specific changes (Maliet et al., 2019; Quintero et al., 2024). In particular, Quintero et al. (2024) recently developed the Birth-Death Diffusion (BDD) process, in which diversification rates diffuse along branches according to a geometric Brownian motion process. The BDD model does not integrate a fossil sampling process, which prevents reliably inferring heterogeneous extinction rates. However, it can incorporate information from the fossil record by constraining extinction trajectories independently inferred for the clade (or subclades) from fossil data. The model has now been extended into the Fossilized Birth-Diffusion (FBDD) process (Quintero et al., 2025), which accommodates fossils placed in the phylogeny. Both the BDD and FBDD models have been implemented in the Julia package Tapestree (Quintero and Andréoletti, 2020), with an inference technique called Bayesian data augmentation (see Methods and Tanner and Wong (1987); Maliet and Morlon (2022)). Diversification analyses in Tapestree operate on fixed phylogenies, but Bayesian data augmentation has also been used for full inference of phylogeny and diversification processes (Barido-Sottani and Morlon, 2023; Chabrol et al., 2025). Data augmentation circumvents some mathematical and computational challenges of computing likelihoods of incomplete observed data; it also produces posterior samples of the full diversification histories, i.e. complete phylogenies including unsampled lineages and, in the case of diffusion models, heterogeneous diversification rates evolving along all (observed and unobserved) branches. From this posterior sample of complete phylogenies, one can directly obtain the distribution of estimated diversity-through-time trajectories for a clade, as well as study in detail the patterns of speciation and extinction at the lineage level. For example, using the BDD model, Quintero et al. (2024) observed an imbalanced speciation process in mammals wherein most lineages undergo a progressive decline in speciation rates before going extinct - notably at the K-Pg mass extinction event - while a few lineages preserve a high speciation potential and drive the diversification of their clade. The selective extinction of lineages with low speciation rates could only be captured by taking into account the rates of the unobserved lineages. Similarly, using the FBDD model, Quintero et al. (2025) studied diversity trajectories across 27 radiations and used data-augmented histories to show that diversity expansions are fueled by bursts of rapid speciation, while slowdowns and declines are primarily governed by extinction rates. The current implementation of FBDD aims to correct for a mismatch between the FBD process, which assumes a Poisson sampling of fossils, and paleontological databases, in which morphospecies are often only represented by a single described specimen (Wright et al., 2022). To this end, taxonomical information can be used in the FBDD inference to add fossil occurrences corresponding to the morphospecies included in the phylogeny. Although this reduces the mismatch mentioned above, the model does not incorporate a specific sampling process for occurrences and ignores occurrences that do not belong to included morphospecies.

Building on these recent developments, we introduce the Occurrence Birth-Death Diffusion (OBDD) model, and implement it in Tapestree, together with its simpler homogeneous variant, the Occurrence Birth-Death (OBD) model. These models combine the strengths of their predecessors while tackling both the computational limitations of the previous OBD implementations and the need to accommodate rate heterogeneity across lineages. They extend the analytical framework by accurately incorporating a broader range of fossil records, including those without described and recorded morphological characters or reliable taxonomic information at the subclade level, which are often very numerous. These developments allow us to tackle an important interrogation, namely assessing the relative contribution of phylogenies, various types of fossils, and rate heterogeneity to accurately reconstructing diversification histories. After developing and validating the OBD and OBDD models, we explore this question by applying a range of diversification models, accounting or not for different fossil data types and rate heterogeneity, to a cetacean dataset.

## Materials & Methods

### The Occurrence Birth-Death Diffusion process

We introduce the Occurrence Birth-Death Diffusion (OBDD) model, which combines a total-evidence approach of the fossil record with the flexibility of a branch-specific heterogeneous birth-death model (Fig. 1). To model heterogeneity in diversification rates across the phylogeny, speciation and extinction rates are assumed to evolve according to a geometric Brownian motion (“GBM”), following Quintero et al. (2024). That is, the logarithms of speciation and extinction rates follow a Brownian motion (see Fig. 1) with variance parameters *σ*^2^ and *σ*^2^, respectively. Furthermore, trend parameters 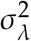 and 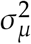 detect consistent rate increases or decreases over time. Specifically, the GBM processes start with initial values *λ*_0_ and *μ*_0_, respectively, and follow:

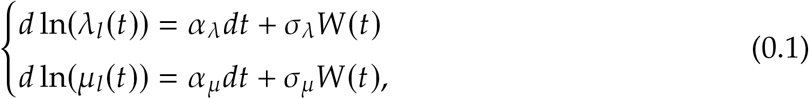

with *λ*_*l*_(*t*), *μ*_*l*_(*t*) being the branch-specific rates for lineage *l* at time *t*, and *W* (*t*) the Wiener process (i.e., standard Brownian motion).

**Figure 1.**
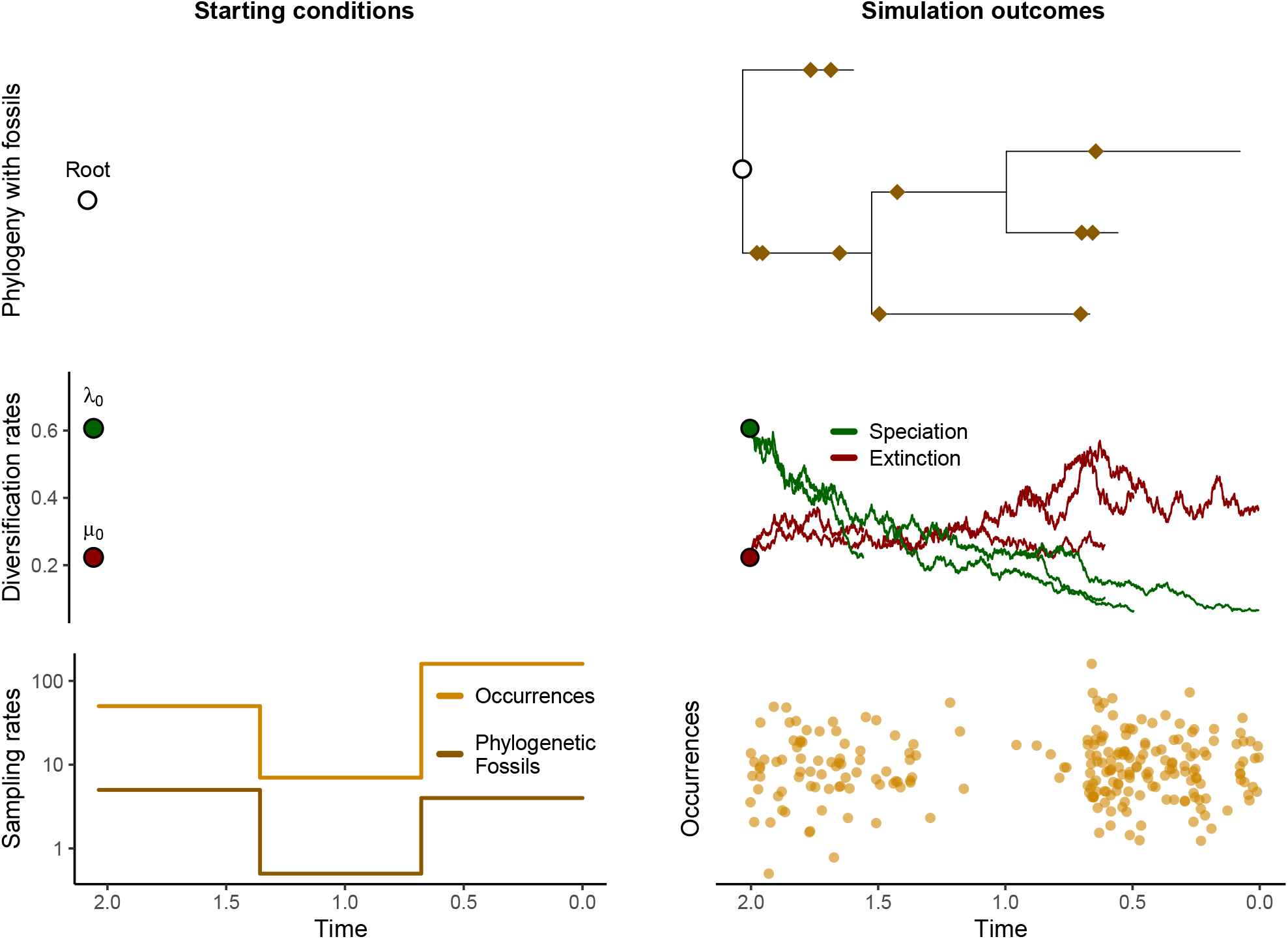
The Occurrence Birth-Death Diffusion process. This figure illustrates a simulation under the OBDD process. One key feature is that speciation and extinction rates are not specified in advance, except for starting values (resp. *λ*_0_ and *μ*_0_), but instead change gradually along each new branch of the phylogeny. The other important feature is that the sampling process produces fossil occurrences that are not placed in the phylogeny, in addition to the standard phylogenetic fossils. Consequently, each simulation will result in both a sample of rate trajectories (right column, middle) and a phylogeny with fossils and a record of occurrences (right column, resp. brown diamonds and orange circles).

Fossils are incorporated into the phylogeny with a fossil sampling rate (*ψ*) following a Poisson point process along branches of the tree (Stadler, 2010; Heath et al., 2014). The integration of fossil occurrences employs a similar approach, with an independent rate of occurrence sampling denoted by *ω* (Vaughan et al., 2019). The difference is that these occurrences are not included in the phylogeny but are instead only recorded as sampling times (as shown in Fig. 1). Given that fossilization and fossil recovery rates are expected to vary over geological time (Dominici et al., 2020), we allow these sampling rates to vary in a piecewise manner at user-defined times (Fig. 1).

### Parameter inference with Bayesian data-augmentation

We implement the OBDD model within the computational framework of the Julia package Tapestree (Bezanson et al., 2017; Quintero and Andréoletti, 2020). Parameter inference uses a Bayesian data augmentation approach (Tanner and Wong, 1987), as described in prior studies on other phylogenetic diversification models (Maliet and Morlon, 2022; Barido-Sottani and Morlon, 2023) and on Birth-Death Diffusion processes in particular (Quintero et al., 2024, 2025). During a Markov Chain Monte Carlo (MCMC) procedure, the data augmentation algorithm involves alternately simulating missing parts of the phylogenetic tree, referred to as augmented branches, as well as all branch-specific diversification rates, and updating the model parameters based on this augmented data. Therefore, a complete phylogeny, combining the reconstructed tree and the augmented branches, with associated diversification rates, is available at each iteration of the MCMC, simplifying the likelihood computation for any set of model parameters (see Appendix A). Data augmentation thus circumvents the difficulty of computing the likelihood of complex models based on the reconstructed phylogeny, which otherwise requires the analytical integration of unobserved components of the process. Furthermore, this iterative process generates a “forest” of posterior augmented trees, with associated diversification rates, which can be used to reconstruct the posterior distribution of species richness, or evolutionary rates, over time.

To model rate heterogeneity across the phylogeny, we discretise all branches at regular intervals, *δt* and approximate the GBM described in the previous section. Gibbs sampling is used for most parameters, leveraging conjugate priors to enhance the efficiency of the inference process, and a Metropolis-Hastings algorithm is used for augmenting unobserved branches (Quintero et al., 2024, 2025). The fossil sampling rate (*ψ*) is inferred similarly to the FBD and FBDD processes, using a conjugate Gamma prior updated based on the number of fossils and the total branch length of the complete augmented phylogenies, as in Quintero et al. (2025). In the following, we detail parts of the inference particular to the OBD and OBDD process, i.e., those involved in the integration of fossil occurrences (sampling rate *ω*).

The MCMC procedure explores the parameter space guided by the likelihood of observed data for current and proposed parameter values. Given that we introduced fossil occurrences, let us consider their likelihood according to our sampling process. On a single branch of length *L*_*i*_ with a constant sampling rate *ω*, the likelihood of observing a specific number of fossil occurrences *k*_*i*_ follows a Poisson distribution 𝒫(*ωL*_*i*_). When *N*_*i*_ lineages exist in parallel, the sum of all sampled occurrences also follows a Poisson process 𝒫(*ωL*_*i*_ *N*_*i*_). We apply this property across *I* intervals delimited by changes in the diversity-through-time curve of the augmented tree: each interval *i* has a constant number of lineages *N*_*i*_ of identical length *L*_*i*_ and contains *k*_*i*_ occurrences. The likelihood of the whole occurrence dataset Ω given the complete augmented tree Ψ and the occurrence sampling rate *ω* is thus the following:

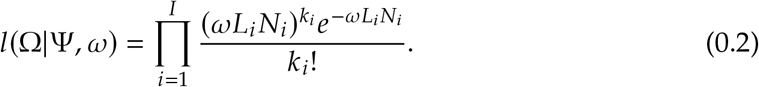

Let us compute the likelihood ratio for a new proposal parameter value *ω*′ relative to the current value *ω*:

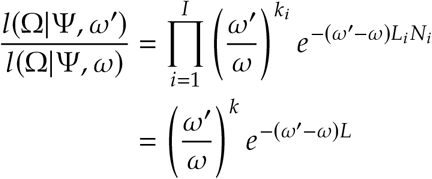

where 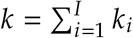 is the total number of occurrences and 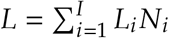 is the cumulative length of all observed and augmented lineages. Note that the exact timing of the occurrences does not affect the likelihood ratio; the process is equivalent to considering a single Poisson process 𝒫(*ωL*) on the full augmented tree. With piecewise-constant sampling rates, the same reasoning applies within each time period. Thus, the posterior distribution of *ω*_*p*_ in a period *p* can be efficiently sampled using a conjugate Gamma prior, Γ(*γ*_1_, *γ*_2_), which is updated based on the observed fossil occurrences *k*_*p*_ and sum of branch lengths *L*_*p*_ during this period:

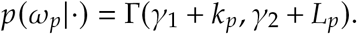

The data augmentation steps involve proposing modifications to the tree, that is, replacing some of the reconstructed branches by subtrees simulated under the process, and accepting or rejecting these proposals based on Hasting ratios. Here, the acceptance ratio used for the inference of the FBDD process (Quintero et al., 2025) is modified to account for the likelihood ratio of the occurrence data given the proposed and current augmented phylogenies. The likelihood of observing the fossil occurrences given an augmented tree depends only on the number of lineages over time, represented by the Diversity-Through-Time (DTT) trajectory. Modifying the augmented tree alters the DTT, which in turn affects the likelihood of the fossil occurrence data. Using the likelihood expression in Equation 0.2 over *J* intervals delimited by changes in the current or proposed DTT, we derive the following likelihood ratio of the proposed tree Ψ′ compared to the current tree Ψ:

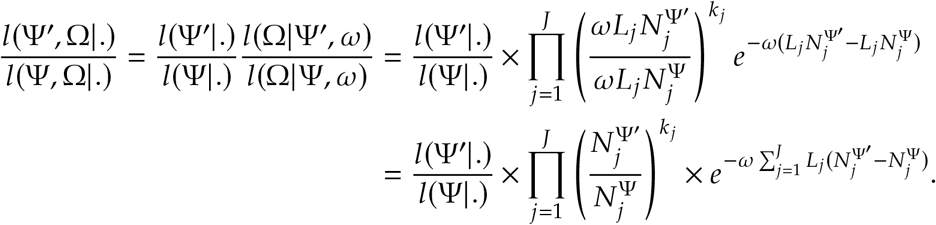

The first part of this ratio is identical to the one used in the FBDD process (see Appendix A), while the added terms ensure that at each iteration the augmented trees reflect the observed fossil occurrence record. In other terms, the greater the number of occurrences included in a given period, the more likely it is that many augmented branches will be accepted in that interval, which will then influence the diversification rate estimates.

The simpler OBD model with constant homogeneous speciation and extinction rates is also implemented in Tapestree.jl (Quintero and Andréoletti, 2020) but as an independent model, extending the constant-rate FBD model (Quintero et al., 2025). The integration of occurrences relies on the same functions for the Gibbs sampling and Metropolis-Hastings steps described above.

### Simulation-based model validation

We validate both the OBD and OBDD models using a procedure called simulation-based calibration, which consists, for a given model, of:

1. choosing prior distributions for the model parameters;
2. simulating many trees using parameter values drawn from these distributions (here, we simulated 1000 trees under the OBD model, and 500 under the OBDD model);
3. for each simulated tree, inferring posterior distributions for all parameters, starting from the prior distributions above;
4. counting the recovery proportion of the true parameter values in posterior credible intervals of increasing widths (from 5% to 95%).

If the recovery proportion matches the widths of all credible intervals, the model is considered well calibrated.

For the OBD model, the initial speciation and extinction rates were drawn from a Gamma(1.0, 1.0) distribution, with a time of origin set to 2.0 units. For the OBDD model, the initial speciation and extinction rates were drawn from Gamma(1.5, 1.0) distributions, trend parameters *α*_*λ*_ and *α*_*μ*_ from Normal(0.0, 0.1) distributions, and diffusion parameters *σ*_*λ*_ and *σ*_*μ*_ from InverseGamma(5.0, 0.5) distributions. The time of origin was set to 1.0 units. These distributions were selected to produce phylogenies of moderate size for computational efficiency, while covering realistic parameter values (Quintero et al., 2025). Fossil sampling rates *ψ* and fossil occurrence rates *ω* were modelled with piecewise constant rates over two epochs, divided at half the time of origin, and are now denoted as *ψ*_1_, *ψ*_2_, *ω*_1_, *ω*_2_. They were drawn, respectively, from the following distributions: *ψ*_1_, *ψ*_2_ ~ Gamma(1.0, 1.0) and *ω*_1_, *ω*_2_ ~ Gamma(1.0, 2.0) distributions for OBD and *ψ*_1_, *ψ*_2_ ~ Gamma(2.0, 1.0) and *ω*_1_, *ω*_2_ ~ Gamma(2.0, 5.0) for OBDD, reproducing the expected higher number of fossil occurrences compared to phylogenetic fossils. MCMC inferences were run for 500.000 and 2.000.000 iterations, respectively. We recorded the total running time of each MCMC inference, in order to analyse how runtime scales with phylogeny size and the inclusion of fossils.

### Case study: Cetacea

We used the Cetaceans as a case study to explore the effect on the inference of diversification histories of i) the inclusion of fossils, in the phylogeny or as simple occurrences, and ii) accounting for rate heterogeneity. We used the cetacean phylogeny from Lloyd and Slater (2021), which contains 89 extant species and 405 fossils (representing distinct morphotaxa). This phylogeny was reconstructed through a meta-analytic approach to combine previous datasets into a metatree that was then used to set constrains on a timetree inference under an FBD tree prior. More specifically, we used five phylogenies drawn from the set of posterior phylogenies provided in the paper (identified as the “Safe” posterior) as replicates for each analysis. In addition, we used the 966 fossil occurrences from the Paleobiology Database (McClennen, 2024) used in Andréoletti et al. (2022), and we sampled their age uniformly in their age uncertainty range at the beginning of each replicate. Their temporal distributions over the five replicates are presented in Appendix C.

We fitted eight diversification models, all implemented in Tapestree.jl:

- the constant-rate Birth-Death, Fossilized Birth-Death, Occurrence Birth-Death, and Occurrence Birth-Death without fossils models (BD, FBD, OBD, and OBD-noF);
- the Birth-Death Diffusion, Fossilized Birth-Death Diffusion, Occurrence Birth-Death Diffusion, and Occurrence Birth-Death Diffusion without fossils models (BDD, FBDD, OBDD, and OBDD-noF).

For the BD and BDD models, we removed fossils from the phylogenies, as well as the stem branch, which is also a product of adding fossils. For the OBD and OBDD models, we added fossil occurrences. For the OBD-noF and OBDD-noF models, we removed fossils from the phylogeny but included occurrences to assess the relative contribution of fossils within and outside of the phylogeny. We kept the stem branch, as it can be inferred in empirical datasets using occurrences. For models including fossilization (i.e., all models except the BD and BDD models), we accounted for known differences in fossil availability across geological periods by considering piecewise constant temporal variations in fossilization rates (*ψ* for phylogenetic fossils and *ω* for occurrences), enabling rate changes at the start and end of the Early Oligocene (Rupelian), Early Miocene (Aquitalian), and Late Miocene (Messinian) periods of well-established low preservation (Marx et al., 2016; Dominici et al., 2020), as in Andréoletti et al. (2022). We thus inferred seven fossil sampling rates and seven occurrence sampling rates for the OBDD model. Table 1 details the data, the number of parameters, and the MCMC inference efficiency of each of these models on the cetacean dataset. Specific features of the diversification history of cetaceans (see Discussion) required us to run the model for a greater number of iterations and therefore longer than in Figure 4 to ensure good convergence. MCMC runs that were still ongoing after 100 days were interrupted. Convergence was evaluated using Tracer (Rambaut et al., 2018) and the R library *coda* (Plummer et al., 2006), with the average Effective Sample Sizes (ESS) reported in Table 1. This evaluation focuses on parameter convergence rather than augmented tree convergence, as current phylogenetic methods for assessing phylogenetic convergence are limited to trees with known tips. Nevertheless, the two should be closely related in the data augmentation procedure.

**Table 1.** Summary of the different diversification models and corresponding analyses on the cetacean dataset. The table shows for each analysis the number of model parameters, characteristics of the cetacean data subset included in the inference - number of tips, fossils and fossil occurrences - and the computational efficiency. Models include homogeneous variants (BD, FBD, OBD, OBD-noF) and diffusion-based variants (BDD, FBDD, OBDD, OBDD-noF); the suffix -noF indicates models run without fossils in the phylogeny, retaining only fossil occurrences. MCMC durations are averaged over 5 replicates. Effective sample sizes (ESS) are averaged over replicates and parameters.

| Model | BD | FBD | OBD | OBD-noF | BDD | FBDD | OBDD | OBDD-noF |
| --- | --- | --- | --- | --- | --- | --- | --- | --- |
| Parameters | 2 | 9 | 16 | 9 | 5 | 13 | 20 | 13 |
| Tips | 89 | 372 | 372 | 89 | 89 | 372 | 372 | 89 |
| Fossils | 0 | 405 | 405 | 0 | 0 | 405 | 405 | 0 |
| Occurrences | 0 | 0 | 966 | 966 | 0 | 0 | 966 | 966 |
| Burn-in | 50K | 100K | 100K | 100K | 50K | 50K | 50K | 50K |
| Iterations | 2M | 2M | 2M | 50M | 10M | 10M | 10M | 20M |
| Duration (days) | 0.01 | 0.07 | 0.85 | 2.9 | 7.7 | 100 | 100 | 80 |
| Average ESS | 1300 | 1800 | 1400 | 740 | 810 | 1900 | 1800 | 310 |

## Results

### Validating the OBD and OBDD implementations

Our simulation-based calibration analysis of the Occurrence Birth-Death (OBD) and Occurrence Birth-Death Diffusion (OBDD) models demonstrate a well-calibrated implementation for all parameters (Fig. 2 and 3). For the OBD model, the recovery proportions of the true parameter values consistently match the credible interval widths (2a) and the estimation errors decrease with increasing tree size (2b). The OBDD model is similarly well calibrated for all parameters (Fig. 3a), and the estimation errors for both speciation and extinction rates diminish as the tree size grows (Fig. 3b). Overall, the validation confirms that both models perform reliably in recovering true diversification parameters from simulated data.

**Figure 2.**
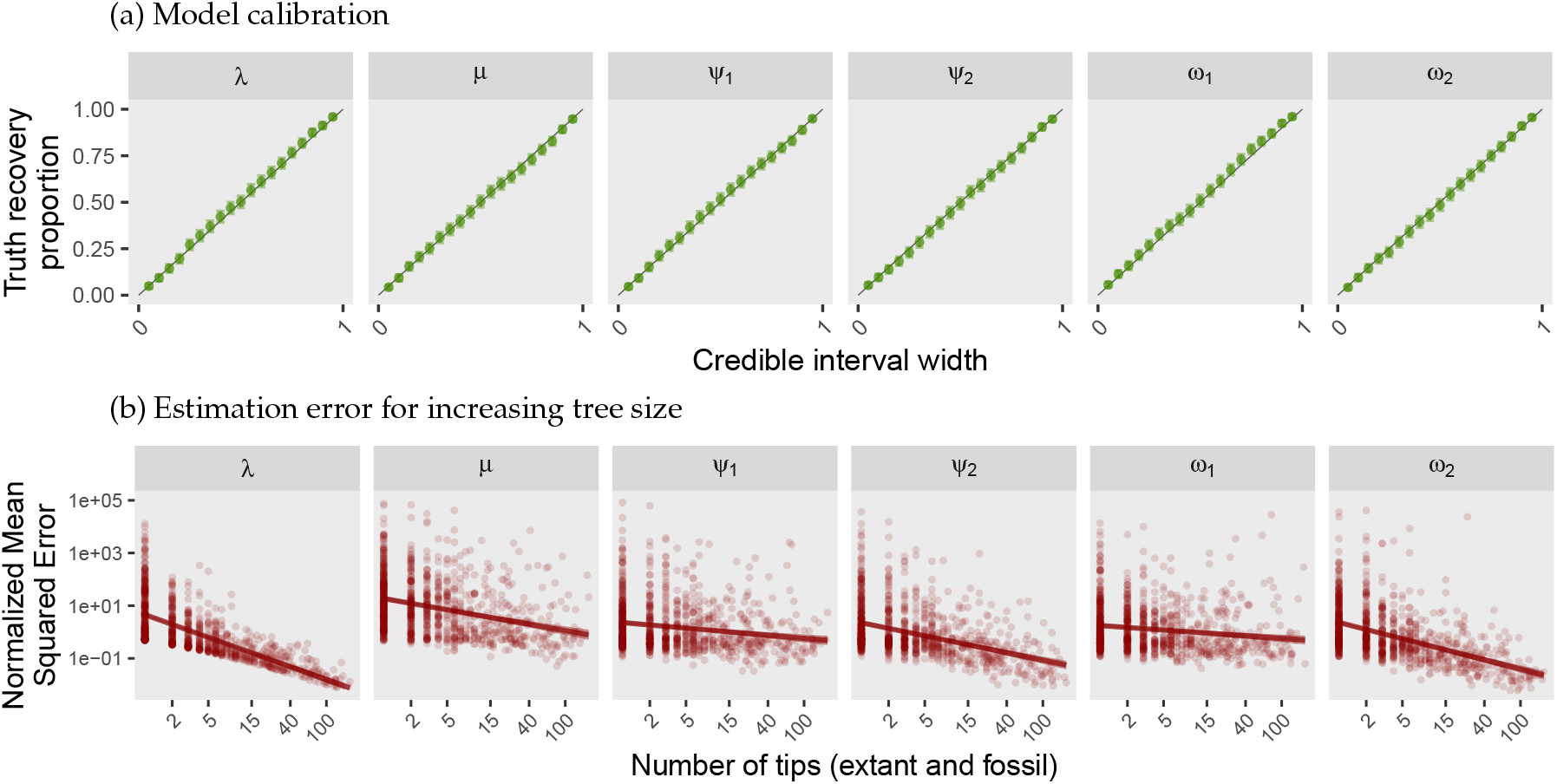
Simulation-based validation of the OBD model.

**Figure 3.**
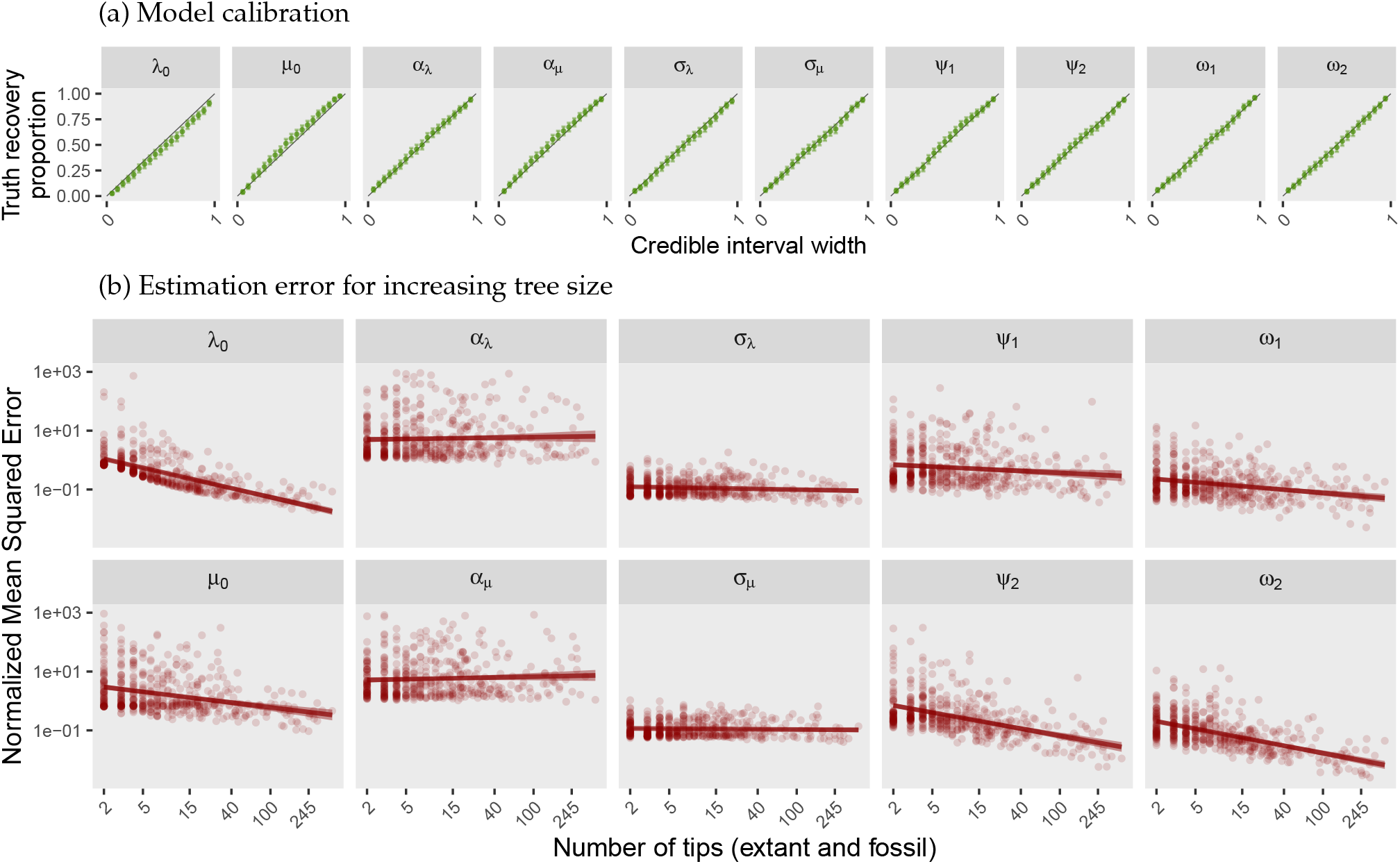
Simulation-based validation of the OBDD model.

Regarding the influence of tree size on computation time, Figure 4 shows that, for datasets generated under the OBDD model and for a fixed number of MCMC iterations, our inference should take an extra day for each additional 150 tips in the reconstructed phylogeny (including fossil tips). The Generalised Linear Model (GLM) and Generalised Additive Model (GAM) regressions (see Appendix B) revealed no influence of the number of fossils or occurrences on the runtime, as apparent in Figure 4, indicating that the code has been adequately optimised for extensive fossil integration.

**Figure 4.**
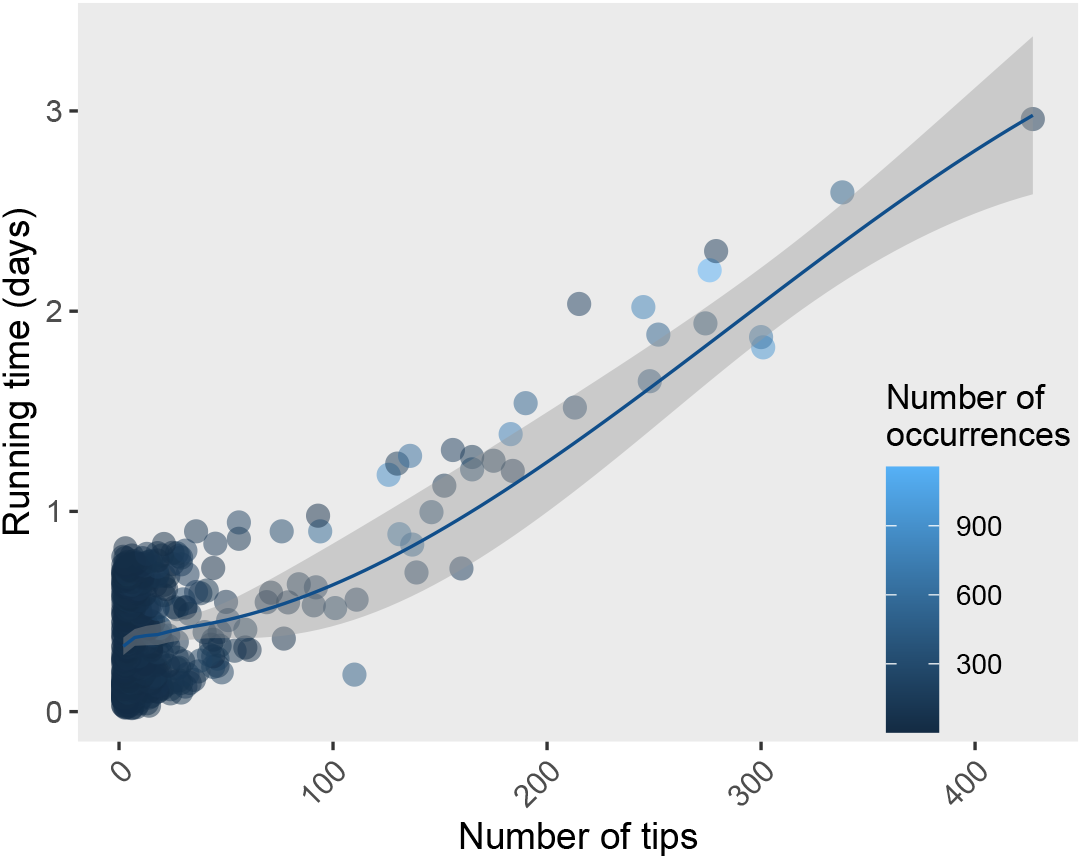
Scaling of MCMC runtime with tree size for the OBDD model. Each point corresponds to one of the datasets used for the simulation-based model validation procedure, from which we recorded the number of tips in the reconstructed phylogeny (including fossil tips) and the runtime for the set number of iterations. The colour gradient indicates the number of simulated occurrences.

In Appendix D, we further explore the drivers of estimation error for each parameter using the same simulated datasets. We observe that the error reduction for several parameters is mostly driven by their corresponding data: extant tree size for the initial speciation and extinction rate, number of fossils for the fossil sampling rates, and number of occurrences for the occurrence sampling rate. The drift and diffusion parameters are generally more difficult to infer (see Fig. 3b), and no significant predictor influence was detected. The interaction between fossils 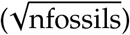 and occurrences 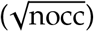 also has an influence on the estimation error, which was positive for most parameters, although only significant for *ψ*_2_ and *ω*_2_. The positive effect reflects that as the number of fossils included in the phylogeny increases, the information provided by additional occurrences decreases, and vice versa.

### Assessing the role of fossils in estimating the diversification history of Cetaceans

Consistent with previous results on mammals (Quintero et al., 2024), we found that models without fossils infer overly simplistic species richness trajectories, close to the observed lineages through time (LTTs) of the reconstructed phylogenies (Fig. 5a and 5e). These models fail to capture the rise and fall trajectory of cetacean diversity expected from paleontological records over the Cenozoic (Morlon et al., 2011; Marx and Fordyce, 2015; Quintero et al., 2025). In models that include only fossil occurrences but exclude phylogenetic fossils (OBD-noF, OBDD-noF, Fig. 5b and 5f), we recover diversity estimates of similar magnitude to the inferred richness trajectories by models with fossils fully integrated into the phylogeny (Fig. 5c–d and 5g–h). However, the uncertainty of the estimate is much higher.

**Figure 5.**
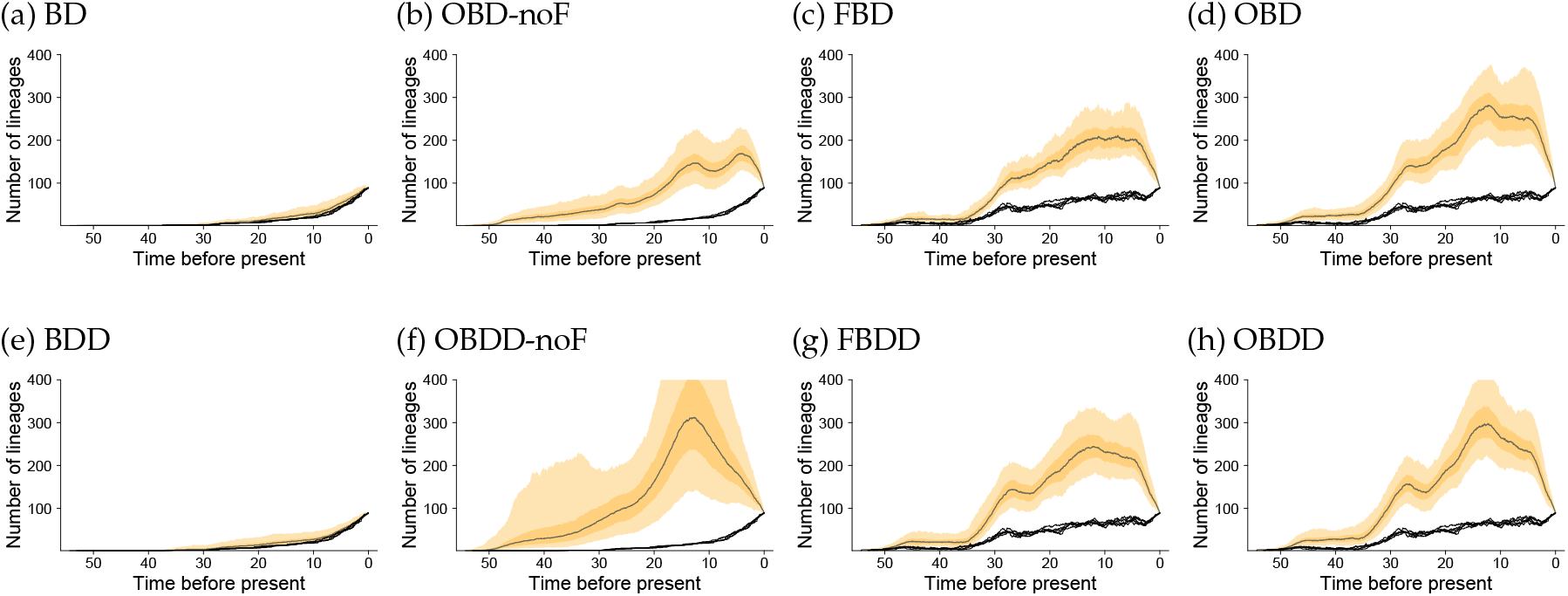
Inferred species richness over time across models. Each panels shows the inferred number of cetacean lineages over time with a given model, conditioned on the reconstructed tree and recorded occurrences. The gray line is the median value across the augmented trees in the combined MCMC samples from the 5 replicate analyses. The 5 overlapping black lines represent the lineage-through-time curves from the reconstructed phylogeny of each replicate. Dark and light yellow areas represent 50% and 95% credible intervals.

The diffusion models (BDD, OBDD-noF, FBDD, OBDD) infer richness trajectories that are qualitatively similar to their homogeneous counterparts, albeit with higher maximum richness levels (especially for OBDD-noF). However, looking at the average diversification rates over time, Fig. 6g and 6h demonstrate that, even though the overall richness patterns are comparable, the incorporation of rate diffusion detects clear variations in the rate trajectories. Both the FBDD and OBDD models infer a rise of speciation rates up to 35 million years ago, followed by a slow decrease, while extinction rates exhibit three major peaks at approximately 40, 27, and 3 Ma, either matching or most recently exceeding speciation rates. In contrast, the BDD model, as shown in Figure 6e, infers low and nearly constant speciation rates (~0.15 per million species years, MSY), with extinction rates that steadily decline over time and are lower than speciation rates for most of the group’s history. Constant rate models infer speciation and extinction rates that are close to averaged heterogeneous rates (see panels 6a to 6d). The BD model (6a) infers the lowest rates and higher speciation than extinction, while the models with phylogenetic fossils and/or occurrences infer negligible net-diversification rates.

**Figure 6.**
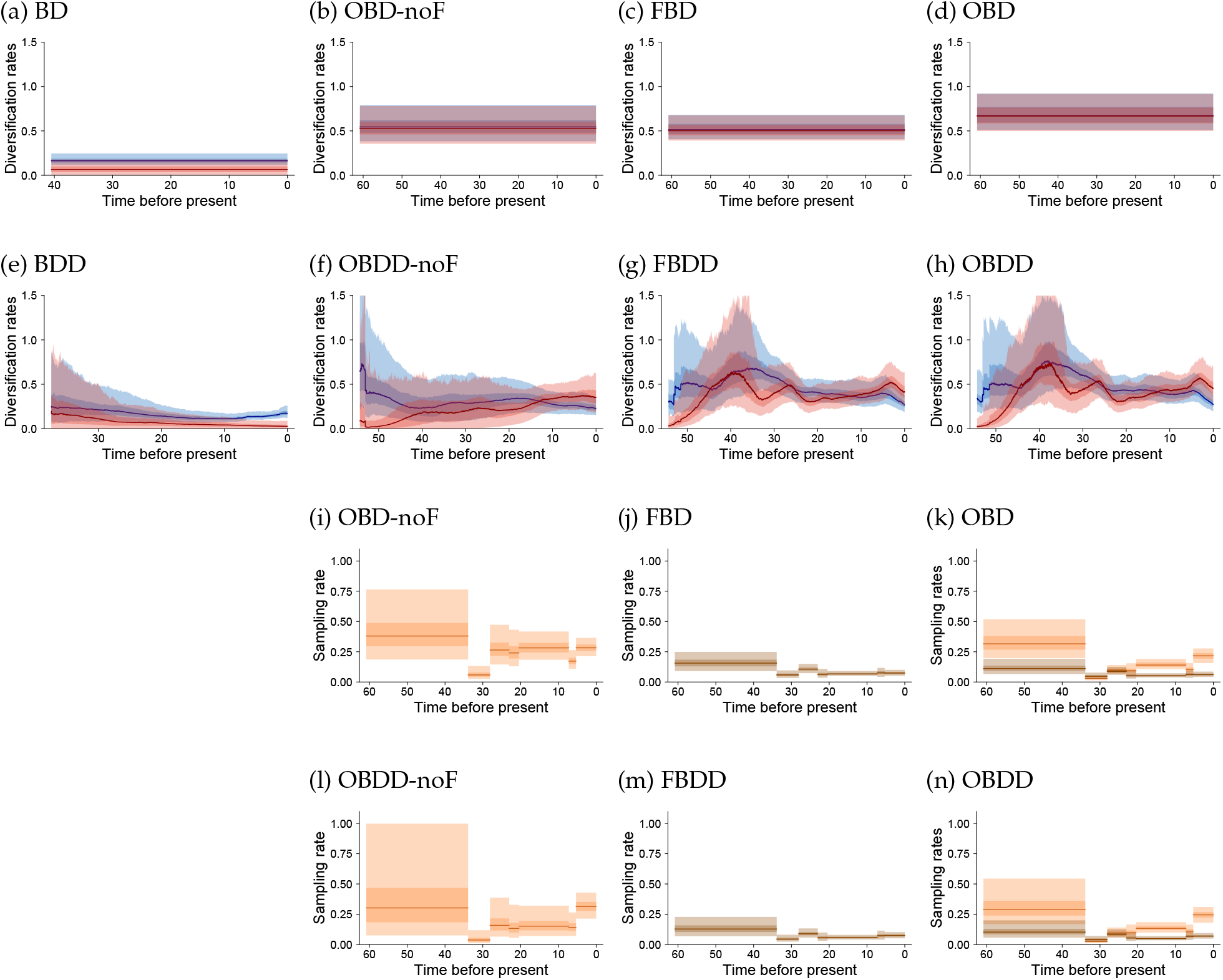
Inferred cetacean diversification and sampling rates across diversification models. Panels a-h show inferred speciation (blue) and extinction (red) rates over time. The line is the median value averaged at each given time over all the branches, observed or augmented, in the combined MCMC samples of the 5 replicate analyses. Panels i-n show inferred piecewise-constant sampling rates of phylogenetic fossils (brown) and occurrences (orange) over time, when applicable. For all panels, dark and light shaded areas represent 50% and 95% credible intervals.

The reconstructed diversity trajectories for constant-rate models with fossils (Fig. 5b–d) may seem surprising given the estimated diversification rates (Fig. 6b–d), as hump-shaped diversity curves are not expected under constant net diversification rates. In fact, the diversity curves reconstructed here should not be interpreted as expected diversity curves under the inferred models, as in Etienne et al. (2011); Billaud et al. (2019), but as curves conditioned on the observed data. The past diversity estimates are indeed reconstructed from the distribution of augmented complete phylogenies produced by the data augmentation procedure, which is conditioned on the reconstructed phylogeny and fossil occurrences. This distribution is therefore equivalent to the diversity curve conditioned on the data for the homogeneous OBD model described in Manceau et al. (2021), whose algorithm was implemented by Andréoletti et al. (2022) in RevBayes.

The remaining panels in Figure 6 show sampling rates for the models with fossils. Occurrence sampling rates are higher than those of phylogenetic fossils, as expected, and the inferred rates in each time period are generally very similar between homogeneous (6j–i) and heterogeneous (6m–l) models. Both sampling rates are reduced in the 3 delimited geological periods of expected low preservation (see Methods), and particularly in the Rupelian (33,9-28,1 Ma).

### Heterogeneous Cetacean diversification

The inclusion of heterogeneous rates in diffusion models reveals significant variation in diversification rates between different clades. Based on the phylogeny of extant species, the BDD model infers markedly higher net-diversification rates for the Delphinidae family, up to 0.4 per Million Species Years (MSY), as shown in Figure 7m, compared to other lineages that exhibit rates between −0.1 and 0.2 per MSY. This difference is mainly due to higher speciation rates in this family (panel 7e) rather than reduced extinction.

**Figure 7.**
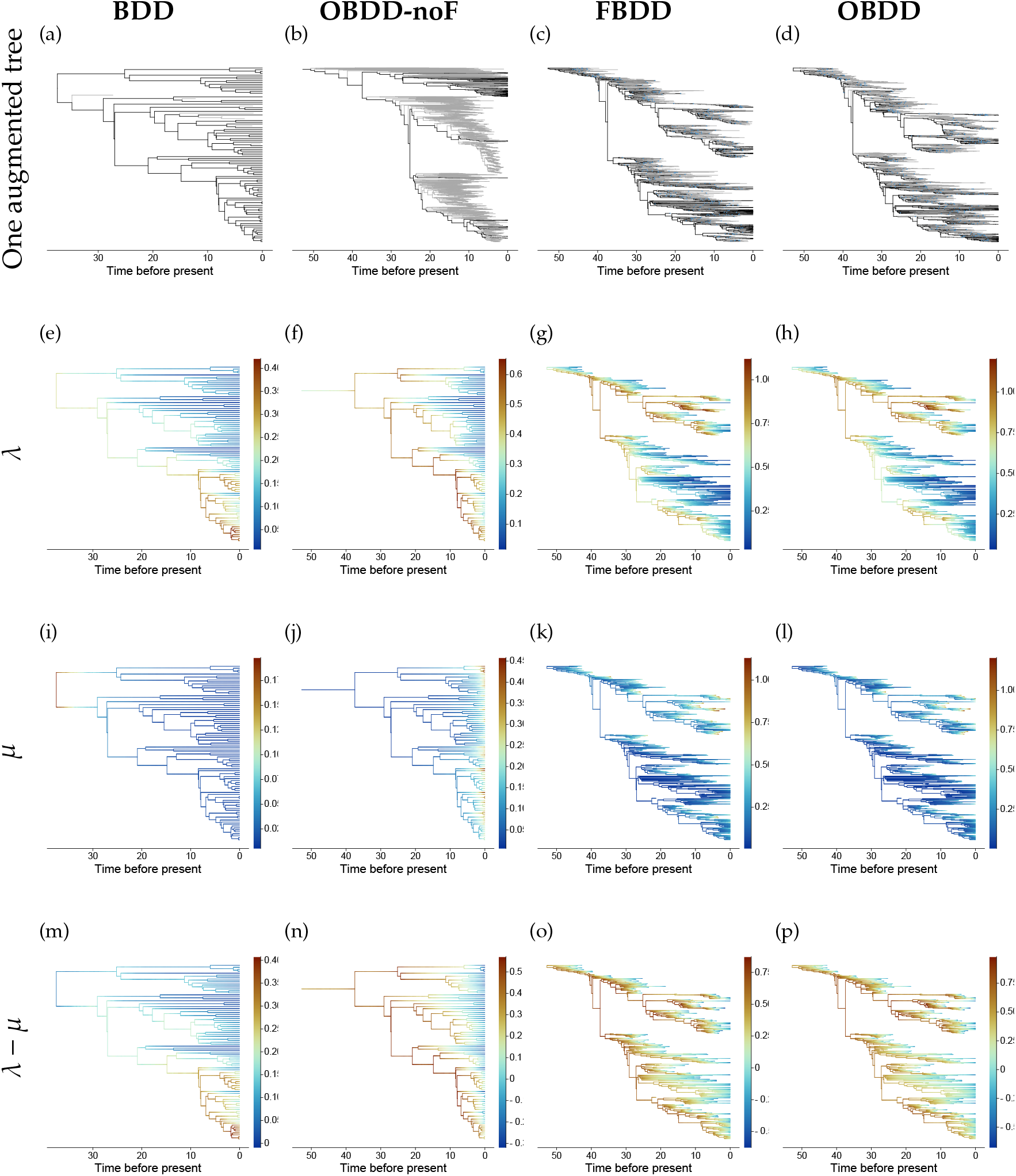
Branch-specific cetacean diversification rates across diffusion models. Panels a-d display samples of complete phylogenies from the MCMC posteriors, with reconstructed branches in black and augmented branches in gray. The remaining panels show the median posterior rates on the reconstructed phylogeny, with branch-specific speciation, extinction and net-diversification rates on the second (e-h), third (i-l) and fourth (m-p) row, respectively.

The diffusion models with phylogenetic fossils, including FBDD and OBDD, capture even stronger levels of rate heterogeneity across cetacean lineages (panels 7o and 7p), with most clades exhibiting high net-diversification rates at their origins, followed by a progressive decline, except for those that gave rise to modern cetacean families. This trend is due, in most individual lineages, both to a strong decline in speciation rate (panels 7f–h) and to an increase in extinction (panels 7j–l).

## Discussion

### Fossils are key to understand the past under BDD models

Our results demonstrate the importance of incorporating fossil evidence to accurately reconstruct diversification histories. The two models we used here that rely solely on phylogenies (BD and BDD) drastically underestimate the historical diversity of cetaceans reflected in the fossil record. Since these models are well calibrated, we assume that the discrepancy arises from model misspecification, specifically imposing constant and homogeneous diversification rates for the BD model and geometric Brownian motion dynamics for the BDD model. The integration of fossil evidence likely mitigates the impact of these misspecifications. Morlon et al. (2011); Mazet et al. (2023) were able to recover a humped-shaped diversity pattern from a phylogeny of extant cetacean species, suggesting that the model they used, which allowed large shifts in speciation and extinction dynamics at the base of major families, better represents the true dynamics of cetaceans. However, making such a priori hypotheses could in turn be the source of model misspecification in other clades. Instead, the flexibility offered by models such as the BDD, if complemented with fossil information, can recover accurate dynamics and be more immune to misspecification issues. In the case of cetaceans, BDD models enhanced with fossil data were able to recover diversification shifts at the base of families and boom-bust diversity dynamics, even though such dynamics are rather unlikely under the BDD process. This is consistent with the findings of Quintero et al. (2024) showing that BDD models can recover clade-wide shifts in diversification rates.

The integration of fossil occurrences shows the value of these additional data, in particular in the absence of fossils whose phylogenetic placement can be inferred: occurrences alone improve the reconstruction of hidden diversity. However, they are less informative than fossils placed in the phylogeny, especially when many of these fossils are available, as is the case for cetaceans. We expect occurrences to be most valuable for accurately estimating diversification rates in epochs where fossil information from phylogenies is sparse, or in groups that lack such fossils altogether. In some groups, it should be possible to use other data reflecting a clade’s total species richness over time as a type of occurrence that can be used in addition to the phylogeny. These alternative occurrences could include teeth, scales, plant organs (leaves, seeds, pollen), biochemical proxies, ichnofossils (trace fossils), coprolites. Many of these remains can be identified as part of the clade, but cannot be associated to other parts of the whole organism that are used for species identification (e.g. the skeleton) and are as such described in parallel parataxonomies that usually cannot be mapped to the clade’s main taxonomy. This fact thus prevents them from being incorporated in the phylogeny in an FBD analysis, but not as occurrences in an OBD analysis, as long as they can be unambiguously attributed to the clade of interest.

Importantly, the fossil record is inherently incomplete, limiting our direct observation of past diversity. This incompleteness poses a significant challenge for methods that rely solely on the observed fossil record (Silvestro et al., 2014). In contrast, the phylogenetic modeling framework, including all models presented here, capitalizes on phylogenetic relationships to infer the existence of unsampled lineages. This represents one of the advantages of the total-evidence approach to reconstruct past diversification histories.

### Heterogeneous rates give us a refined view of a clade’s evolution

Interestingly, rate heterogeneity appears to have a limited influence on the estimated past diversity, which is already well constrained by fossils and occurrences. This is notable given that heterogeneous models were compared to constant-rate homogeneous models, whereas time-varying models may capture additional temporal dynamics. The inclusion of heterogeneity in the diffusion models (BDD, FBDD, OBDD) is thus most valuable for flexibly capturing variations in diversification rates at various scales, and using them to generate hypotheses on the evolutionary pressures, such as environmental changes or clade-specific factors, that may have influenced the diversification of a clade. These hypotheses can then be tested using models specifically designed for this purpose (Morlon et al., 2024).

At the individual lineage level, we observe a general decline in net-diversification rates (similar to the “insidious loss of species macroevolutionary fitness” described in Quintero et al. (2025)), except for a few lineages that maintain their speciation potential and give rise to most of the subsequent diversity. This selection effect matches the pattern of imbalanced speciation pulses observed in Quintero et al. (2024), as well as the theoretical predictions under a model of evolutionary tempo by Budd and Mann (2025).

### The diversification history of cetaceans based on their complete fossil record

The occurrence-based models (OBD and OBDD) recover a hump-shaped pattern, in line with previous fossil and molecular studies (Morlon et al., 2011), but with higher diversity. In particular, they reproduce the rapid decline in the past 5 Myr that has previously been linked to global climate cooling and the Northern Hemisphere glaciation (Andréoletti et al., 2022). This pattern broadly matches the one obtained at the genus level in Andréoletti et al. (2022), although it differs in including a short decline in the late Oligocene (25 Ma) and a maximum attained in the Mid-Miocene (12 Ma) instead of the Pliocene (around 4.5 Ma).

Looking at the average rate trajectories over time, especially the ones inferred with the FBDD and OBDD models which incorporate the most information, it appears that extinction is the main driver of variations in diversification trends on the scales of millions of years, although it is concomitant with a long-term decline in speciation rates that together explain the decline of cetaceans. The second spike of extinction around 27 million years ago corresponds to the extinction of many archaeocetes, the stem cetacean lineages, soon after the emergence of the two modern groups, Mysticeti and Odontoceti. These diversification rate trajectories are consistent with the trends inferred by Quintero et al. (2025) using a variant of the FBDD model. Their analysis incorporates additional occurrences to inform the fossil sampling rate in order to correct for the common failure to include multiple fossils of the same morphospecies, but this only includes occurrences of morphospecies present in the phylogeny. Despite these similar trends, our estimated rates are approximately 50% higher overall than those reported by Quintero et al. (2025), and the lineage diversity in our FBDD and OBDD results is nearly double. This discrepancy likely arises from our approach of distinguishing the processes for sampling phylogenetic fossils on one side and fossil occurrences on the other. In Quintero et al. (2025), including occurrences as additional sampled ancestors increases the sampling rate in the tree, which in turn decreases the estimated diversity, speciation rates, and extinction rates.

Considering variations among cetacean lineages, most long-lived species tend to see their net-diversification rate progressively decline, including many modern species. An exception is the Delphinidae family, which diversified rapidly in the past 10 Myr and displays higher net diversification rates close to the present. This likely reflects a combination of late Miocene–Pleistocene oceanographic reorganizations (closure of Tethys and the Central American Seaway, fluctuating seaway connections around southern Africa) and large sea-level oscillations, which repeatedly created opportunities for allopatric splits across oceanic basins, together with some delphinid traits (broad diets, social foraging) that allowed them to thrive under shifting, nutrient-rich environments driven by upwelling (Gowans et al., 2007; Steeman et al., 2009; do Amaral et al., 2018). The findings align with Barido-Sottani and Morlon (2023), which identified the highest speciation rates in the Delphinidae family using ClaDS, another heterogeneous diversification model with branch-specific rates. A key methodological difference between the two studies is that Barido-Sottani and Morlon (2023) inferred the phylogeny together with the diversification rates, while we applied here the diffusion models on a fixed reconstructed phylogeny. This phylogeny was obtained using an FBD prior with homogeneous diversification rates, which could generate biases (Barido-Sottani and Morlon, 2023). The convergence of results obtained here versus in Barido-Sottani and Morlon (2023) suggests that the sequential approach (i.e., reconstructing the phylogeny first) can be robust. Still, it would be useful in the future to allow joint reconstruction inference with diffusion models.

## Conclusion

We introduced the Occurrence Birth-Death Diffusion (OBDD) process, a new model that integrates fossil occurrences and accommodates heterogeneous diversification rates across lineages. Our analyses show that fossils are essential for accurately reconstructing diversification histories under diffusion models. Occurrences provide additional refinements that are particularly useful when phylogenetic fossils are sparse. In cetaceans, accounting for rate heterogeneity revealed important variations among lineages and during lineage lifetimes, as well as several periods of increased extinction.

Looking ahead, the OBDD model opens new avenues. Occurrences can often be attributed to specific subclades, and incorporating this information – similar to the use of topological constraints in traditional FBD analyses (Barido-Sottani et al., 2023) – would provide more direct insights into lineage-specific rates. In some cases, occurrences might be identified at the morphospecies level, offering even finer resolution (Silvestro et al., 2014; Stadler et al., 2018). Additionally, model accuracy may be enhanced by coupling the classical fossil sampling rate with the occurrence sampling rate instead of inferring them independently, as these rates are expected to partially co-vary. These fossil sampling rates could also be allowed to evolve gradually along branches of the phylogeny, capturing variations in sampling across species and extending existing models with heterogeneous sampling processes (Barido-Sottani and Morlon, 2025). These developments augur an exciting trajectory for models that, with ever-greater data integration and flexibility, can gradually uncover more subtle complexities of past evolutionary processes.

## Funding Acknowledgment

This work was supported by the École Normale Supérieure de Paris (doctoral fellowship to J.A.).

## Acknowledgment

The authors thank the members of the Morlon team for their feedback on an earlier version of this manuscript.

## Data Availability Statement

The code and data for reproducing the analyses are available at https://github.com/Jeremy-Andreoletti/OBDD.

## Authors’ contributions

J.A. and H.M. designed the project. I.Q. contributed the initial BDD and FBDD codes. J.A. wrote the OBDD code, performed the analyses, and wrote the first draft of the paper. All authors discussed the research and contributed to the writing and editing of the article.

## APPENDIX

### A. Likelihood of the complete tree

Following Quintero et al. (2025) for the FBDD model, we approximate the likelihood of a complete tree Ψ under the OBDD model by generating augmented discrete GBM paths along branches of the complete trees. Using the same notations for consistency, we set a small time step *δ* = *t*_*i*+1_ − *t*_*i*_ and divide each branch of the tree into *m* steps such that *m* = ⌊ *t*_*b*_/*δ*⌋ + 1, where *t*_*b*_ represents the edge length of branch *b*. For a branch *b*, which starts at *t*_*s*_ and ends at *t* _*f*_, we sample the data augmented diffusion process at times *t* = {*t*_1_ = *t*_*s*_, *t*_2_ = *t*_1_ + *δ*, …, *t*_*m*+1_ = *t* _*f*_}. We thus obtain the stochastic evolution of speciation Λ_*b*_ = {*λ*_*b*_(*t*_1_), …, *λ*_*b*_(*t*_*m*+1_)} and of extinction M_*b*_ = {*μ*_*b*_(*t*_1_), …, *μ*_*b*_(*t*_*m*+1_)} in a discrete time grid. Thereafter *λ*_*l*_(*t*_*i*_) is simplified as *λ*_*i*_ (and thus *λ*_*l*_(*t*_*b*_) = *λ*_*b*_) and *μ*_*l*_(*t*_*i*_) as *μ*_*i*_.

The combined likelihood for the complete tree Ψ (including internal branches Ψ_*I*_ and extinct branches 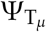) and record of occurrences Ω given the OBDD model parameters as well as the branch-specific speciation and extinction rates, Λ and M respectively, is the following:

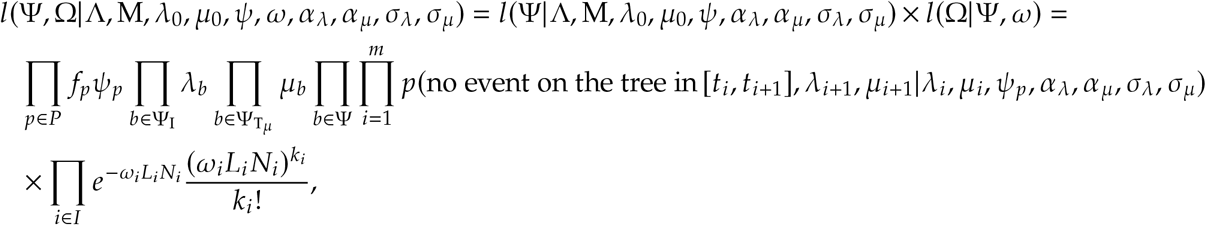

with *f*_*p*_ the number of phylogenetic fossils in the period *p* and *k*_*i*_ the number of occurrences in each interval *i* delimited both by the *P* periods of sampling rate change and the times of change in the diversity-through-time curve, such that in each interval of length *L*_*i*_ the occurrence sampling rate *ω*_*i*_ and the number of lineages *N*_*i*_ are constant.

As in Quintero et al. (2025), the probability that no event happens on the tree during time *δ* = *t*_*i*+1_ − *t*_*i*_ and that the rates starting at *λ*_*i*_ and *μ*_*i*_ end up at *λ*_*i*+1_ and *μ*_*i*+1_ can be approximated as:

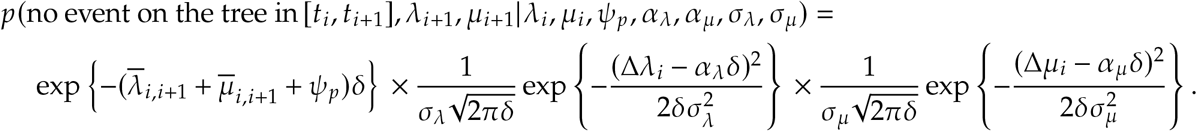

### B. MCMC runtime

The running time for MCMC inferences under the OBDD model was measured across the 500 simulated datasets from the validation analysis and plotted as a function of the number of tips (including fossil tips). The results are shown in Figure 4.

We then applied a Generalized Additive Model for Location, Scale, and Shape (GAMLSS), using the *gamlss* package (Rigby and Stasinopoulos, 2005; R Core Team, 2018), to model the running time as a function of the number of extant tips, the number of fossils, and the number of fossil occurrences. The GAMLSS framework allows us to model both the mean and variance structures separately, thus dealing with heteroscedasticity. We used a Gamma distribution for the response variable (running time), which is appropriate for modelling positive continuous data. The mean structure was modelled using a penalized B-spline (P-spline) for the number of extant tips and linear terms for the number of fossils and fossil occurrences. Additionally, the scale (variance) of the model was allowed to depend on the number of extant tips. Only trees with at least 15 tips were retained in order to obtain normally distributed residuals and correct diagnostic plots.

Here is the resulting summary:

~~~
> summary(gamlss_model)
******************************************************************
Family: c(“GA”, “Gamma”)
Call: gamlss(formula = running_time ~ pb(ntipsextant) + nfossils +
  nocc, sigma.formula = ~ntipsextant, family = GA(mu.link = “identity”,
  sigma.link = “log”), data = subset(parameters_OBDD, par == “lambda” & ntips > 15))
Fitting method: RS()
------------------------------------------------------------------
Mu link function: identity
Mu Coefficients:
                  Estimate Std. Error t value Pr(>|t|)
(Intercept)       0.2627590 0.0287025 9.155 8.15e-16 ***
pb(ntipsextant)   0.0056901 0.0002565 22.183 < 2e-16 ***
nfossils         −0.0009767 0.0005983 −1.632   0.105
nocc              0.0001322 0.0001147  1.152   0.251
---
Signif. codes: 0 ‘***’ 0.001 ‘**’ 0.01 ‘*’ 0.05 ‘.’ 0.1 ‘ ‘ 1
------------------------------------------------------------------
Sigma link function: log
Sigma Coefficients:
                Estimate Std. Error t value Pr(>|t|)
(Intercept) −0.2930249 0.0756514 −3.873 0.000165 ***
ntipsextant −0.0075807 0.0007572 −10.011 < 2e-16 ***
******************************************************************
~~~

While the number of extant tips has a significant non-linear effect on the running time, we found no significant relationship between the running time and the number of fossils or number of fossil occurrences. These results indicate that tree size mostly determines the computational burden and that including fossils have a negligible impact.

### C. Cetacean occurrences over time

Occurrence ages are sampled uniformly within their age uncertainty ranges at the beginning of each analysis. Figure C.1 represents their temporal distribution for the 5 random seeds that were used.

**Figure C.1.**
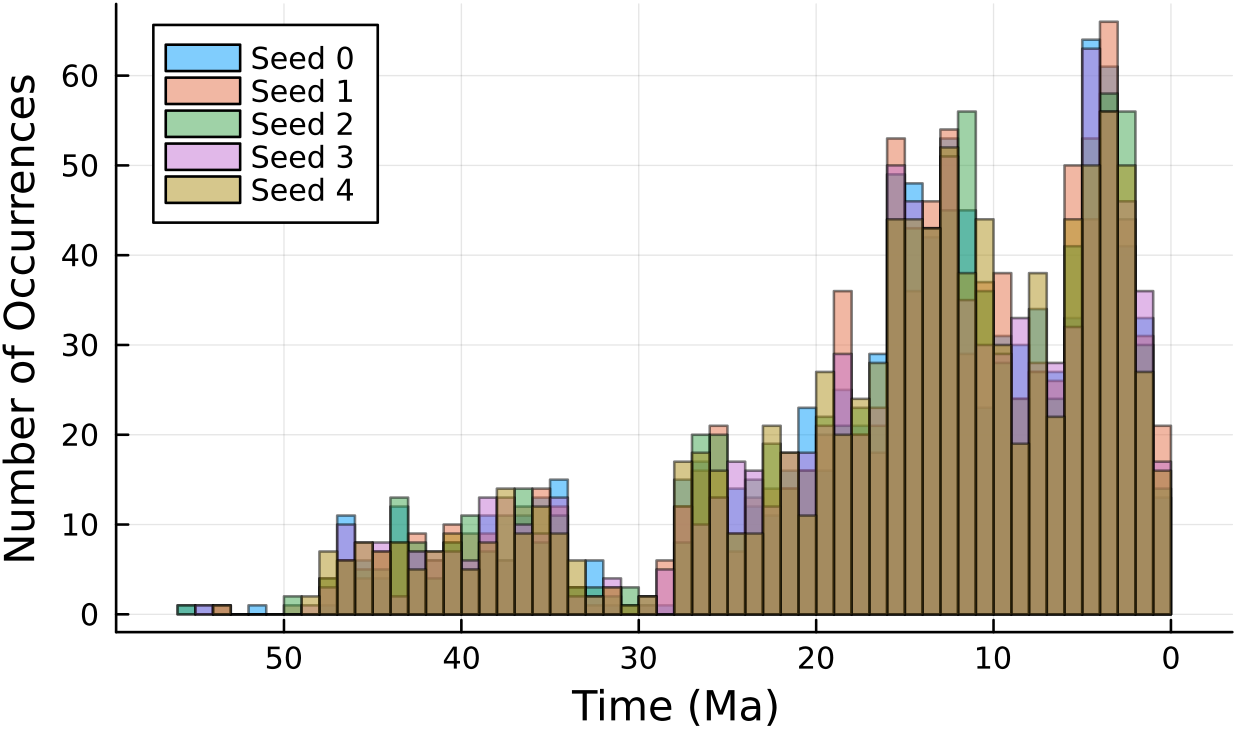
Temporal distribution of cetacean occurrences with age uncertainty. These histograms illustrate the temporal distribution of cetacean occurrences over the Cenozoic. Results are overlaid for the five different sampling seeds used to uniformly sample within the age uncertainty ranges of the fossil occurrences.

### D. Drivers of estimation error

To explore the drivers of estimation error in the OBDD model parameters, we employed a generalized linear model (GLM) with the *stats* package in R (R Core Team, 2018) to predict the Normalized Mean Squared Error (NMSE) for each parameter based on some characteristics of the simulated datasets from the validation analysis. These predictors are the number of extant tips, the number of fossils, the number of fossil occurrences, and the interaction between fossils and occurrences. We extracted estimated coefficients from these models and calculated the predicted mean influence of each predictor on the estimation error.

The analysis covered the following OBDD model parameters:

- *λ*_0_: Initial speciation rate.
- *μ*_0_: Initial extinction rate.
- *α*_*λ*_: Speciation rate trend.
- *α*_*μ*_: Extinction rate trend.
- *σ*_*λ*_: Diffusion variance for speciation.
- *σ*_*μ*_: Diffusion variance for extinction.
- *ψ*_1_, *ψ*_2_: Fossil sampling rates in 2 epochs.
- *ω*_1_, *ω*_2_: Fossil occurrence sampling rates in 2 epochs.

The log(NMSE) for each parameter was modelled as a function of the logarithm of the number of extant tips (log(ntipsextant)), the square root of the number of fossils 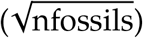, the square root of the number of fossil occurrences 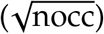, and the interaction between the number of fossils and occurrences 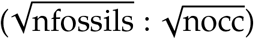. There may be 0 occurrences or fossils, so a logarithmic transformation was not appropriate. For each parameter, we multiplied the coefficients by the average value of the predictors to calculate their predicted influence on estimation error, and display the resulting prediction in Figure D.1.

**Figure D.1.**
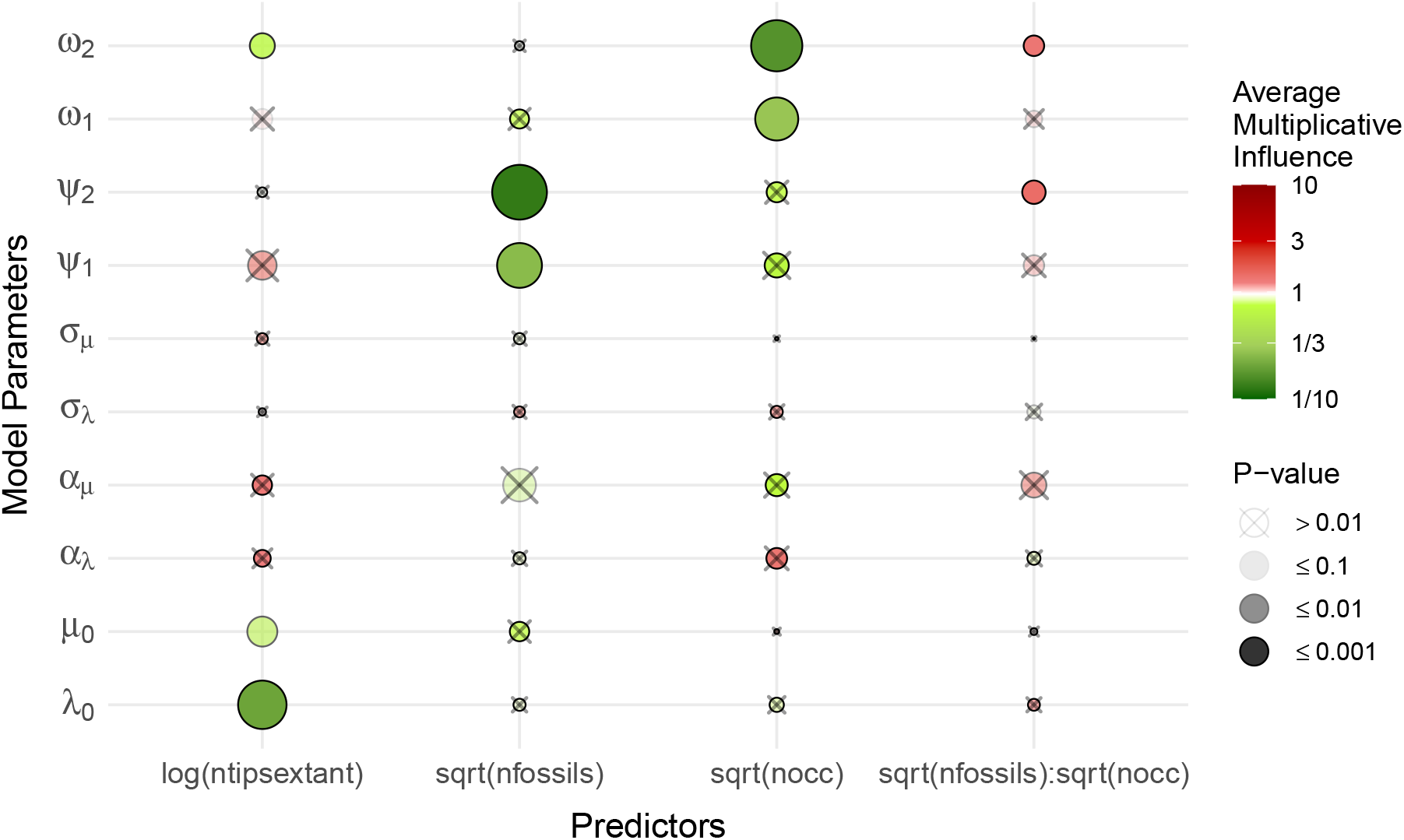
Drivers of estimation error for OBDD parameters. This figure displays the drivers of estimation error, measured as the Normalized Mean Squared Error (NMSE). Each column corresponds to a predictor in a Generalized Linear Model: log of the extant tree size, square root of the number of fossils in the tree, fossils occurrences, and their interaction. Row correspond OBDD model parameters: initial speciation rate (*λ*_0_), initial extinction rate (*μ*_0_), speciation and extinction trends (*α*_*λ*_, *α*_*μ*_), diffusion variances (*σ*_*λ*_, *σ*_*μ*_), and fossil and occurrence sampling rates for two time intervals (*ψ*_1_, *ψ*_2_, *ω*_1_, *ω*_2_). The colour of each circle indicates the average multiplicative influence of a predictor on a specific parameter’s estimation error, and the circle size also reflects the absolute influence in log scale (× 10 has a similar influence as × 1 / 10). The shading of each circle reflects the statistical significance (p-value), with darker circles indicating lower p-values (more significant). Predictors with p-values above 0.01 are crossed.

